# Identification of a new, Rab14-dependent, endo-lysosomal pathway

**DOI:** 10.1101/2021.03.25.436964

**Authors:** Evgeniya Trofimenko, Yuta Homma, Mitsunori Fukuda, Christian Widmann

**Author notes:** Corresponding author: Address correspondence: Christian Widmann, Department of Biomedical Sciences, Rue du Bugnon 7, 1005 Lausanne, Switzerland.

## Abstract

Cells can endocytose material from the surrounding environment. Endocytosis and endosome dynamics are controlled by proteins of the small GTPase Rab family. Several endocytosis pathways have been described (e.g. clathrin-mediated endocytosis, macropinocytosis, CLIC/GEEC pathway). Besides possible recycling routes to the plasma membrane and various organelles, these pathways all appear to funnel the endocytosed material to Rab5-positive early endosomes that then mature into Rab7-positive late endosomes/lysosomes. By studying the uptake of a series of cell-penetrating peptides (CPPs) used in research and clinic, we have discovered a second endocytic pathway that moves material to late endosomes/lysosomes and that is fully independent of Rab5 and Rab7 but requires the Rab14 protein. This newly identified pathway differs from the conventional Rab5-dependent endocytosis at the stage of vesicle formation already and is not affected by a series of compounds that inhibit the Rab5-dependent pathway. The Rab14-dependent pathway is also used by physiological cationic molecules such as polyamines and homeodomains found in homeoproteins. Rab14 is expressed by the last eukaryotic common ancestor. The Rab14-dependent pathway may therefore correspond to a primordial endosomal pathway taken by cationic cargos.

## Introduction

Endocytosis is a major entry route used by cells to take up a variety of extracellular substances ranging from nutrients, fluid phase material, growth factors, hormones, receptors, cellular penetrating peptides (CPPs), viruses or bacteria. Various forms of endocytosis have been described, the main routes being clathrin-mediated endocytosis, macropinocytosis, and the clathrin-independent carrier/glycosylphosphotidylinositol-anchored protein enriched endocytic compartment (CLIC/GEEC) pathway [reviewed in ^1–5^]. Which form of endocytosis is used and the ultimate fate of the endocytosed material depend on the nature of the substances being taken up by cells.

Endocytic vesicles (endosomes) are formed by membrane invaginations, actin-driven membrane protrusions (in the case of macropinocytosis for example), or ruffling. In the case of clathrin-mediated endocytosis, vesicle formation is triggered through the detection of the endosomal cargo by AP2 adaptor domains and subsequent recruitment of clathrin triskelions ^6, 7^. Several AP2 adaptors bound to the plasma membrane through PI(4,5)P2 are necessary for efficient clathrin binding ^7^. Accumulation of AP2/clathrin complexes (within seconds) at the membrane leads to membrane bending and endocytic vesicle formation ^6–8^.

Endosomes are dynamic structures that undergo fusion and fission events ^9^. Early endosomes mature into multivesicular bodies (MVBs), late endosomes and finally lysosomes, where degradation of the endocytosed material occurs ^10, 11^. The endocytosed material can also be recycled back to the plasma membrane or trafficked towards other cellular compartments ^10–12^.

Each stage of endosomal maturation is meticulously controlled by the sequential recruitment of various endosomal protein and lipids. For example, on the early endosomes, Rab5, activated by its guanine exchange factor (GEF) Rabex-5, controls local generation of PI(3)P by recruiting the Vps34 PI3 kinase. This in turn leads to recruitment of EEA1 via its capacity to bind PI(3)P through its FYVE domain. EEA1 can also directly interact with the active GTP-bound form of Rab5. The ability of EEA1 to bind simultaneously Rab5 and PI(3)P on separate vesicles makes it a tethering protein that contributes to endosomal fusion ^13^. Vps34 knock-out in mammalian cells leads to enlarged early endosomes and interruption of the progression of endocytosed cargo to lysosomes ^14^. Vesicle maturation proceeds through the recruitment of the Mon1-Ccz1 complex that interacts with Rab5 and PI(3)P. The Mon1-Ccz1 complex has a GEF activity towards Rab7 that leads to the activation of this small GTPase on endosomes [reviewed in ^11, 15^]. Concomitantly, Rab5 GTPase-activating protein (GAP) turns off Rab5 and promotes release of the latter from early endosomes. Hence, Rab5 and Rab7 regulate essential steps in the endocytic pathway that moves endocytosed material to lysosome. The Rab5/Rab7-controlled endocytic pathway is currently the only molecularly characterized route taken by endocytosed material that end up in lysosomes ^2, 5, 15^.

In this study we show that endocytosed CPPs, homeoproteins, and polyamines, follow a newly discovered endosomal pathway towards lysosomes that requires Rab14 but not Rab5 or Rab7. Endocytosis of CPPs is also unaffected by phosphoinositide 3-kinase (PI3K) inhibitors or various pharmacological agents known to inhibit the uptake of classical cargos such as transferrin and dextran. This work therefore defines a second independent endocytic maturation pathway that moves endocytosed material to lysosomes.

## Results

### CPPs employ unconventional endocytosis

CPPs can be used for intracellular transport of bioactive cargo into cells ^16–28^. Various non-exclusive mechanisms of CPP endocytosis have been proposed ^16–23, 27, 29–31^. However, there is no consensus and clarity regarding the precise nature of the endosomal pathway used by CPPs and its underlying mechanisms. CPPs additionally enter cells through direct translocation via water pores that are formed as a consequence of membrane megapolarization induced by the CPP themselves and the activity of potassium channels ^32^. Direct translocation can be inhibited through plasma membrane depolarization or invalidation of specific potassium channels (e.g. KCNN4 in HeLa cells), without affecting endocytosis of CPPs ^32^, transferrin ^33^ or vesicular stomatitis virus (VSV) ^33^. Here, we took advantage of KCNN4 knockout HeLa cells to study specifically endocytosis in the absence of possible confounding effects mediated by CPP direct translocation. To investigate the endocytic pathway employed by CPPs, we phenotypically characterized CPP containing vesicles (Figure S1A-B) for the presence of early (Rab5 and EEA1) and late (Rab7 and Lamp1) endosomal markers. We selected five most commonly used CPPs in research and in clinic (TAT, R9, Penetratin, MAP and Transportan) as well as TAT-RasGAP317-326, a prototypical TAT-cargo complex ^32, 34–44^. Pulse-chase experiments (Figure 1A-B and S1B) demonstrated colocalization of transferrin, EGF and dextran with EEA1, Rab5A and Rab5B at early time points, and Rab7 and Lamp1 at later time points. These results are consistent with previous knowledge that these molecules enter cells through clathrin-mediated endocytosis (transferrin and EGF) and macropinocytosis (dextran). To ensure that ectopic expression of endosomal markers does not interfere with normal endocytosis, we compared the pattern of EEA1-positive and Lamp1-positive vesicle, which we found to be qualitatively similar in control cells and in cells expressing ectopic GFP-tagged versions of these markers (Figure S1C). Moreover, ectopic expression of the tagged EEA1 and Lamp1 constructs did not alter the kinetics of transferrin colocalization with EEA1-or Lamp1-positive vesicles (Figure S1D). These results indicate that ectopic expression of fluorescent endosomal markers, in live cells in particular, does not appear to affect endocytic processes. CPPs were found in EEA1-and Lamp1-positive vesicles at early and late time points, respectively (Figure 1A-B and S1A) but surprisingly only a minority of CPP-containing vesicles were positive for Rab5 and Rab7 (Figure 1A-B). Even though the selected CPPs have different physico-chemical properties they all carry positive charges within their sequence and appear to be found in the same endocytic vesicles (Figure S1E).

**Figure 1.**
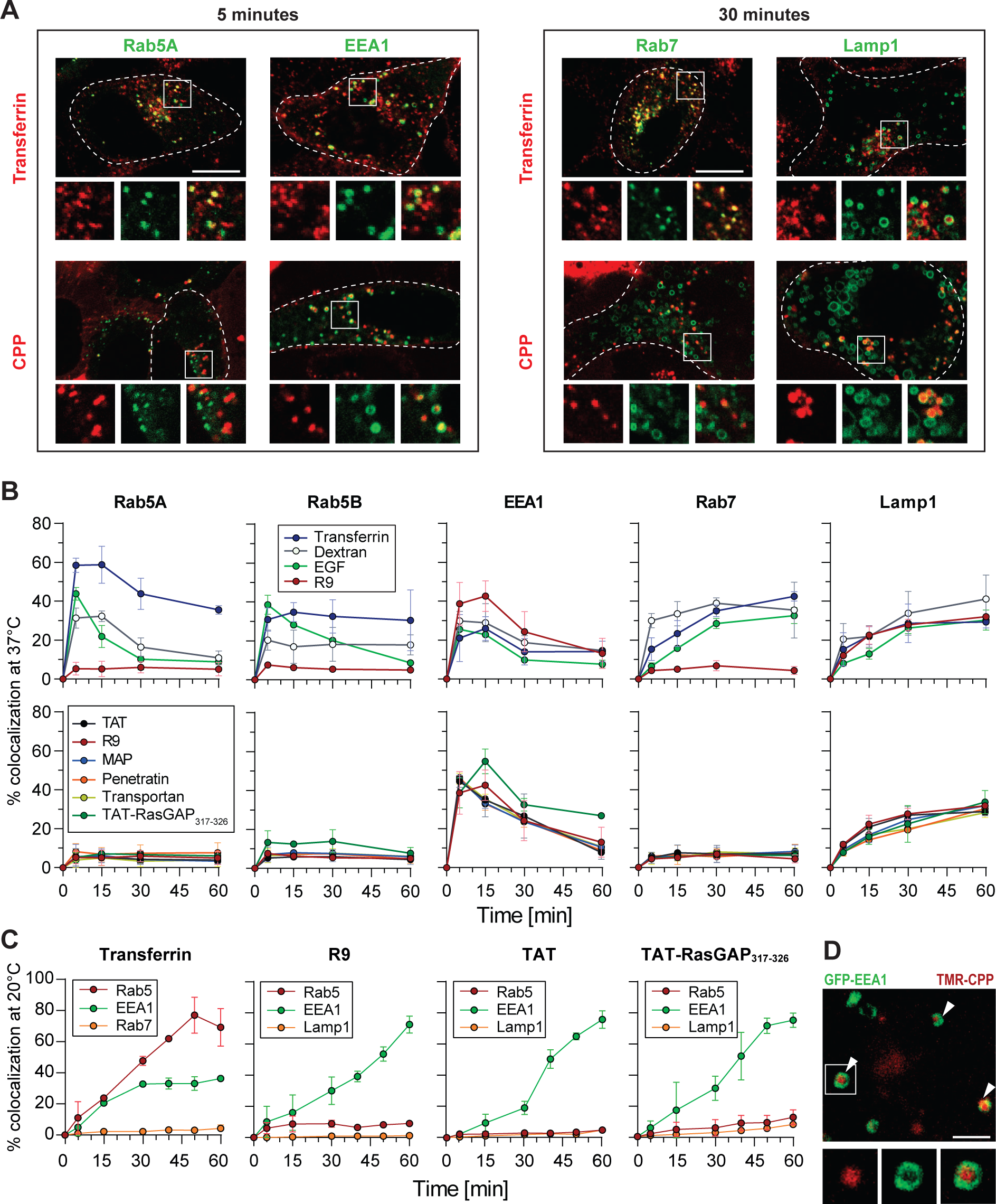
CPPs follow a Rab5-independent endocytic pathway. **(A-D)** HeLa KCNN4 knock-out cells ectopically expressing the fluorescently-tagged early (Rab5A, Rab5B and EEA1) or late (Rab7 and Lamp1) endosomal markers were incubated with 20 μg/ml transferrin, 0.2 mg/ml dextran 10 kDa, 2 μg/ml EGF or with 40 μM of CPPs linked or not to a cargo for 5 minutes, then washed and imaged at the indicated time points by confocal microscopy (see Figure S1B for experimental setup). All experiments were performed on live cells at 37°C unless indicated otherwise. **(A)** Representative confocal images of HeLa KCNN4 knock-out cells, expressing GFP-tagged endosomal markers, incubated with AlexaFluor568-transferrin or TMR-TAT. Images were acquired at 5 or 30 minutes after the addition of endosomal material. Scale bar: 10 μm. **(B-C)** Quantitation of colocalization between the indicated fluorescent material and fluorescently-tagged early and late endosomal markers. Colocalization analysis was performed as described in Methods and Figure S1A. The data correspond to mean ± SD of three independent experiments. In panel B, the data obtained with R9 are shown, for comparison, both on the top graphs and the bottom graphs. In panel C, cells were incubated at 20°C to delay endosomal maturation. **(D)** Representative confocal, Airyscan acquired, high-resolution images of cells ectopically expressing GFP-EEA1 incubated with 40 μM TMR-TAT for 5 minutes. Scale bar: 2 μm.

To rule out that association of Rab5 with CPP-containing vesicles could be transient, we performed experiments at 20°C such that endosomal maturation is considerably slowed down, as can be observed for transferrin (Figure 1C, left). However, CPP-positive endosomes remained mostly Rab5-negative (Figure 1C). Additionally, Rab5- and Rab7-positive CPP-containing vesicles were only marginally detected in the continuous presence of the CPPs despite extensive colocalization with EEA1 (Figure S1F). High resolution confocal images showed that TAT was found inside EEA1-positive vesicles already a few minutes after being added to cells (Figure 1D) confirming that CPPs are indeed located in EEA1-positive endosomes. In addition, TAT-RasGAP317-326 did not interfere with the progression of transferrin through early and late endosomes (Figure S1G) and did not lead to generation of aberrant endosomes bearing EEA1 and Lamp1 at the same time (Figure S1H). This indicates that CPPs do not reprogram the manner by which cells take up material through classical endocytosis. Lack of colocalization with Rab5A and Rab7 was observed previously for tryptophane/arigine-rich peptide WRAP linked to siRNA ^45^. Additionally, based on visual inspection of representative images in literature, R8 and TAT also colocalize only partially with Rab5A ^46^.

We then used pharmacological agents such as EIPA ^47^, IPA3 ^48, 49^, ML7 ^50, 51^, Jasplakinolide ^52^ and Cytochalasin D ^53, 54^ that block distinct steps of endocytic vesicle formation and maturation, such as actin filament polymerization, Na^+^/H^+^ exchange inhibition, as well as macropinosome formation and closure, to determine whether these steps are parts of the CPP endocytic pathway. We observed no effect of these inhibitors on the internalization of TAT-RasGAP317-326 in contrast to what was seen for dextran uptake (Figure S2A-B). Additionally, as opposed to transferrin internalization, the early stages of CPP endocytosis were dynamin-independent (Figure S2C).

Lipids such as phosphoinositides (PIs) that can be phosphorylated at positions 3, 4, or 5 of the inositol ring, represent another type of markers of endocytosis. For example, PI(4,5)P2 is enriched in the plasma membrane, whereas PI(3)P and PI(3,5)P2 are enriched in early and late endosomes, respectively, and participate in their formation [reviewed in ^55–59^]. In conventional endocytosis, EEA1 is recruited to early endosomes through interactions with Rab5 and PI(3)P ^60–62^. The latter is produced by Vps34, a PI3-kinase that is also recruited by Rab5. We therefore, used pan-PI3K inhibitors (wortmannin and LY294002) and assessed the colocalization between selected endosomal material and EEA1 or Lamp1. Our data show that, in the presence of LY294002, transferrin and dextran endosomal maturation and progression was halted, consistent with observations reported in the literature ^14, 63^ (Figure 2A and S2D). However, colocalization between CPPs and EEA1 or Lamp1 was not affected (Figure 2A). Furthermore, depletion of PI(3,4)P2, enriched on the plasma membrane ^59^, blocks the maturation of clathrin-coated vesicles prior to their fission from the plasma membrane ^64^, and prevents macropinosome closure ^65, 66^. As these events precede Rab5 recruitment, we next determined whether the presence of wortmannin, which at high micromolar concentrations, besides PI(3)P depletion, additionally inhibits PI4-kinases ^67, 68^ and leads to depletion of PI(3,4,5)P3 ^69^ and PI(3,4)P2 ^70, 71^, would have an effect on CPP endocytosis. Indeed, wortmannin treatment almost fully inhibited the endocytosis of transferrin and dextran, as opposed to the tested CPPs, where the number of vesicles was either only marginally reduced (R9) or remained unchanged (TAT) (Figure 2B).

**Figure 2.**
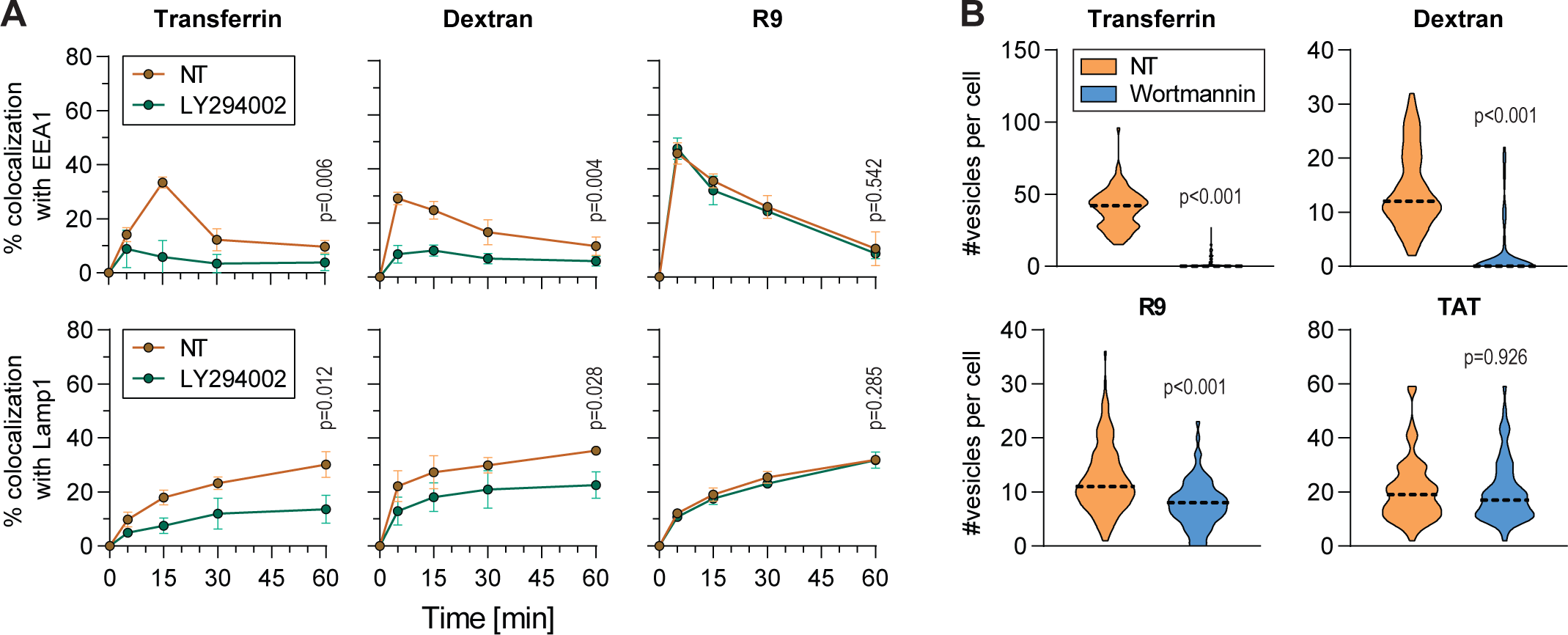
CPP endocytosis does not require PI(3)P-kinase-like enzymes. **(A)** Colocalization quantitation between the indicated endosomal material and endosomal markers in the presence or in the absence of LY294002, a pan-PI3-kinase inhibitor. HeLa KCNN4 knock-out cells, ectopically expressing GFP-tagged EEA1 or Lamp1, were incubated with 20 μg/ml AlexaFluor568-Transferrin, 0.2 mg/ml TMR-Dextran 10 kDa or 40 μM TMR-R9 for a pulse of 5 minutes. Cells were preincubated or not for 30 minutes with 25 μM LY294002, which was still present during the full duration of the experiment. The data correspond to mean ± SD of three independent experiments. The p-values were calculated based on area under the curve (AUC) analysis followed by a t-test. **(B)** Quantitation of the number of endosomal vesicles per cell in the presence or in the absence of wortmannin, a pan-PI3-kinase inhibitor. HeLa KCNN4 knock-out cells were incubated with 20 μg/ml AlexaFluor568-Transferrin, 0.2 mg/ml TMR-Dextran 10 kDa or 40 μM TMR-CPP for a pulse of 5 minutes. Cells were preincubated or not for 30 minutes with 10 μM wortmannin, which was still present during the full duration of the experiment. The number of vesicles positive for the indicated endosomal material were visually calculated based on confocal images, acquired in the middle of the cell. A minimum of 150 cells were quantitated per condition. The results correspond to three independent experiments. The p-values were calculated using paired t-test.

Even though previous studies have shown some contradictory results of CPP internalization in the presence of the endocytic inhibitors used in this study [decreased uptake ^46, 72–74^, no effect ^45, 74–76^, increased uptake ^77^], our data clearly indicate that CPPs enter cells through an uncharacterized endocytic pathway, which differs from classical endocytosis already at the stage of endocytic vesicle formation.

### Rab14 is required for the maturation of CPP-containing endosomes

All previously characterized endocytic pathways appear to converge to Rab5-positive vesicles ^1, 2, 78–87^. In the absence of Rab5, the number of maturing endocytic vesicles is decreased and endocytosis is halted ^62^. To determine whether EEA1 recruitment to CPP-positive vesicles occurs in the absence of Rab5, we took advantage of a knockout Rab library in MDCK cells, which consists of single and multiple knockouts (>50 cell lines) targeting either different protein isoforms or multiple Rab proteins simultaneously ^88^. This library includes a Rab5 conditional knockout cell line, where Rab5A, B and C isoforms have been knocked out and replaced by a Rab5A version that can be degraded through auxin-induced ubiquitination upon addition of indole-3-acetic acid (IAA) ^89^ (Y. Homma et al., manuscript in preparation) (Figure S3A). The recruitment of EEA1 (15 minutes post incubation), as well as Lamp1 (30 minutes post incubation) to TAT-containing vesicles was not affected by the absence of Rab5 (Figure 3 and S3B). Furthermore, inhibiting Rab5 with dominant-negative constructs (Rab5A S34N, Rab5B S34N or Rab5C S35N) in HeLa cells, while reducing the percentage of EEA1-positive transferrin- and dextran-containing vesicles, as expected, had no significant effect on the percentage of EEA1-positive (Figure S3C-D) or Lamp1-positive (Figure S3E) CPP-containing endosomes. This set of data argues against a role of Rab5 isoforms in the maturation of CPP-containing endosomes and for EEA1 recruitment on these endosomes.

**Figure 3.**
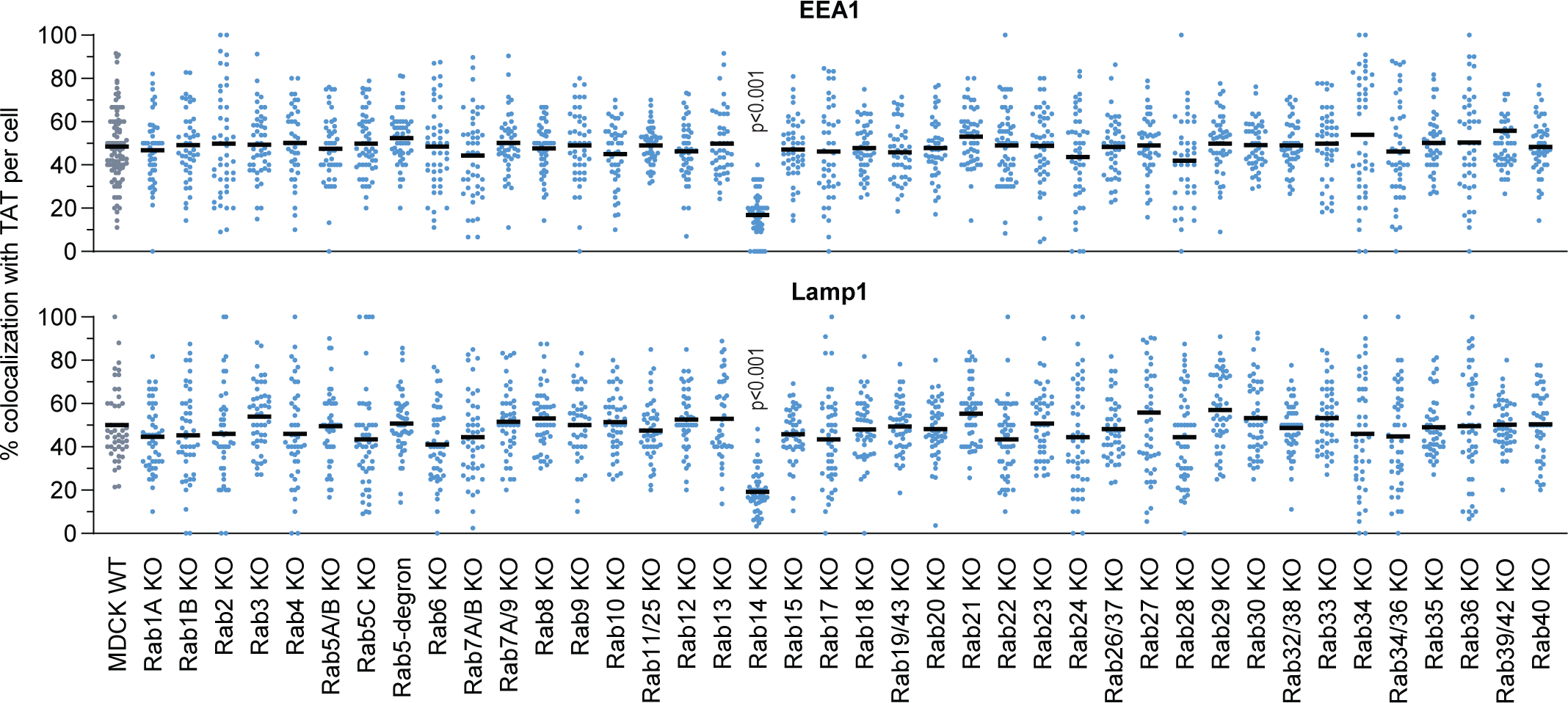
CPP endosomal maturation is Rab14-dependent. Quantitation of colocalization between TMR-TAT and GFP-EEA1 (15 minutes, top) or Lamp1-GFP (30 minutes, bottom) in a pulse chase experiment in MDCK-II wild-type cells and the indicated Rab knock-outs. A minimum of 50 cells were quantitated per condition. Statistical analysis was performed with ANOVA multiple comparison to wild-type condition with Dunett’s correction. Only significant p-values are shown on the Figure. Rab5-degron cells were treated with 1 μg/ml doxycycline and 500 μM IAA for 48 hours to induce degradation of degron-tagged Rab5A in the Rab5B/C knockout cell line.

Using a candidate-based approach and the Blastp database, we identified a subset of candidate proteins with amino acid sequence similarity to Rab5. Using GPS Protein (http://gpsprot.org/navigator.php?q=8411), a database of protein-protein interactions, we restricted the pool of these Rab5-like proteins to those that have the potential to interact with EEA1. We identified three top candidates using this approach: Rab14, Rab22 and Rab31 (also known as Rab22B). Additionally, Rab21 bears sequence similarity to Rab5 and colocalizes with early endosomal markers (Rab5 and EEA1 ^90–92^). Rab14 colocalizes with early endosomal markers ^93–95^, but not late endosomal markers ^93, 95, 96^ (Figure S4) and has so far been shown to be involved in endosomal recycling ^97^, endosome-endosome fusion, and MHC class I cross presentation ^96^. Rab14 and Rab22 are both involved in endosome to Golgi trafficking ^93, 95, 98^, and Rab31 plays a role in Golgi-endosome directional transport ^99^. Rab22 can directly interact with EEA1 ^98, 100^, supporting the hypothesis that it could function in a manner similar to Rab5. Our data showed that Rab14, Rab21 and Rab22 colocalize relatively frequently (as opposed to Rab31) with transferrin ^93, 94^, and CPPs, as well as to a lesser extent with dextran (Figure S5A). Furthermore, dominant-negative versions of these proteins (Rab14 S25N, Rab14 N124I, Rab21 T31N, Rab22 S19N) were used to assess the role of the respective Rab proteins in CPP-containing vesicle maturation. Figure S5 shows diminished colocalization between CPPs and EEA1 in cells expressing Rab14 S25N and Rab14 N124I, but not in cells expressing the other dominant negative mutants. Rab14 is therefore a strong candidate that can substitute for Rab5 function in the recruitment of EEA1 to maturing CPP-containing endosomes. The colocalization between transferrin, as well as dextran and EEA1 appears not to be affected in the Rab14 dominant negative background (Figure S5B). Similarly, earlier work has reported that EEA1 colocalization with mannose receptors ^96^ or with EGF ^95^ is not affected when Rab14 is depleted in cells.

We then used a MDCK-II comprehensive Rab knock-out library to test whether colocalization between CPPs and EEA1 or Lamp1 is affected by the depletion of various Rab protein isoforms at 15-and 30-minutes post incubation. The obtained data further support the involvement of Rab14 in the maturation of CPP-containing vesicles (Figure 3). Surprisingly, none of the other tested single or multiple Rab knockouts had an effect on EEA1 or Lamp1 recruitment to CPP-positive endosomes (Figure 3). Taken together our data demonstrates that ensuing their uptake by cells, CPPs follow a Rab5-independent, Rab14-dependent endocytic route.

### Homeoproteins and polyamines use the same endocytic pathway as CPPs

Homeoproteins (HPs) are a family of transcription factors involved in multiple biological processes ^101–104^. Additionally, HPs, such as Engrailed 2 and OTX2, exhibit therapeutic properties ^105–110^. The vast majority of HPs contain a conserved 60 amino-acids domain called the homeodomain (HD) that carries HP internalization and secretion motifs. Interestingly, the HP internalization motifs bear CPP sequences ^101, 103^. We therefore hypothesized that HDs could be endocytosed similarly as CPPs. Through colocalization experiments we determined that there is indeed colocalization between HD-containing endosomes and the EEA1 or Lamp1 markers, and that their colocalization with Rab5A or Rab7A is only marginal (Figure 4A, left). As for CPPs, EEA1 and Lamp1 recruitment to HD-positive endosomes was significantly reduced in cells lacking Rab14 (Figure 4B, left). This indicates that HDs follow a Rab5-independent, Rab14-dependent endosomal pathway. Polyamines are small signaling molecules involved in numerous cellular processes (gene regulation, cell proliferation, cell survival and cell death) ^111–115^. In mammalian cells polyamines enter cells through endocytosis ^116–119^, and possibly also through a polyamine specific transporter ^117^. Polyamine-containing vesicles mature to Lamp1-containing acidic endosomes and are then exported into the cytosol ^119^. The mechanism of polyamines endocytosis has not been described at the molecular level. Similarly, to CPPs and HDs, polyamine-containing vesicles colocalized only marginally with Rab5A and Rab7A markers and their maturation down to Lamp1-containing endosomes was Rab14-dependent (Figure 4A-B, right and S6). These data indicate that physiological molecules such as polyamines, and by extension homeoproteins if they behave like their homeodomains, do not enter cells through the classical Rab5-dependent endocytic route but via a Rab14-dependent pathway.

**Figure 4.**
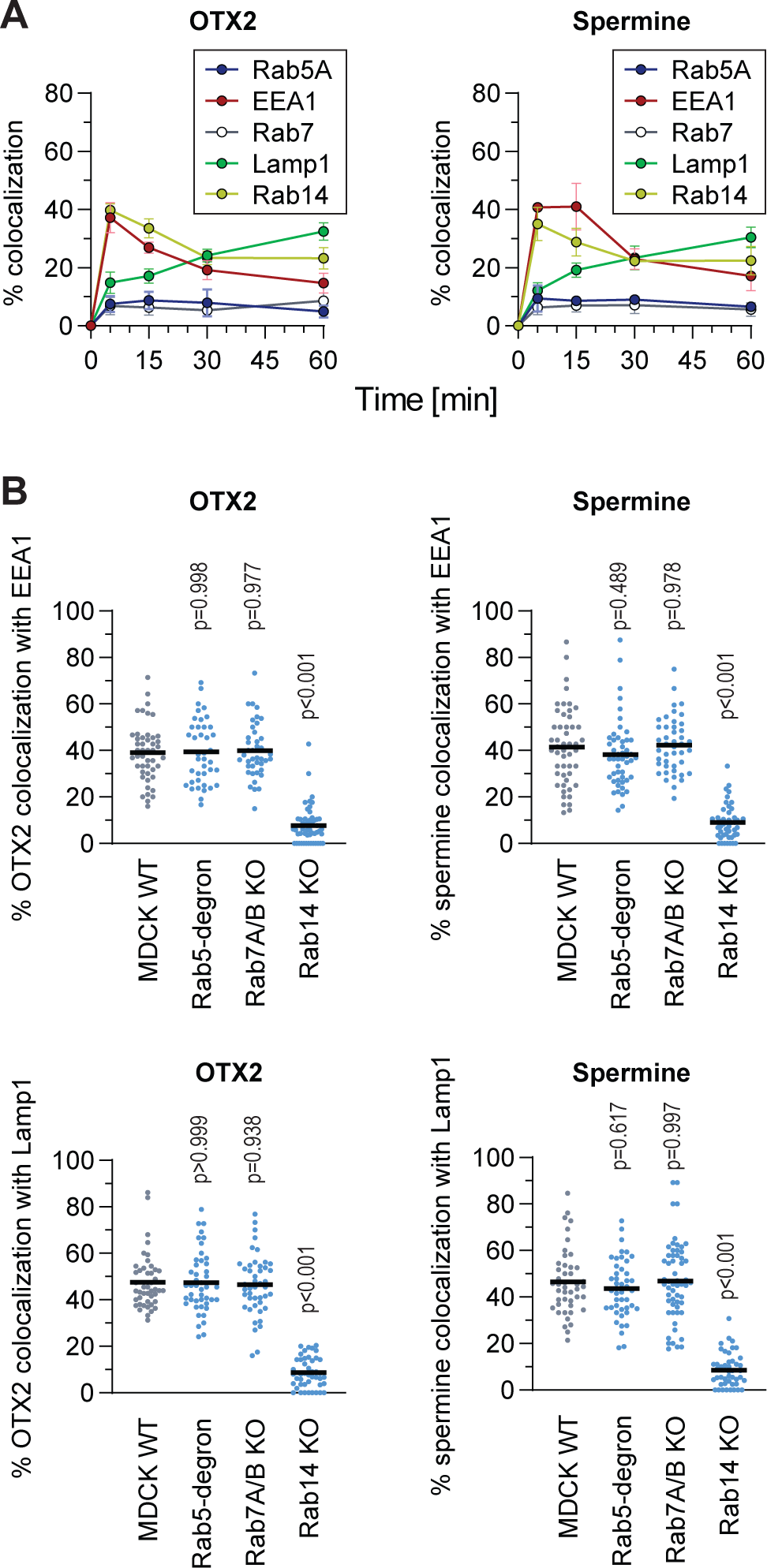
HDs and polyamines are following a Rab5-independent, Rab14-dependent endocytic route. **(A)** Colocalization quantitation between endosomal markers and HD or polyamine. HeLa KCNN4 knock-out cells, ectopically expressing the indicated endosomal markers, were subjected to a 5-minute pulse incubation with fluorescently labelled OTX2 HD (10 μM) or spermine (5 μM). Quantitation assessment was based on confocal images. The results correspond to mean ± SD of three independent experiments. **(B)** Colocalization quantitation between fluorescent versions of OTX2-HD (10 μM) or spermine (5 μM) with EEA1 (top) or Lamp1 (bottom) in MDCK-II wild-type and the indicated Rab knockouts. Cells were incubated for 5 minutes, then washed and confocal images were acquired at 15- and 30-minutes post incubation for early and late endosomal markers, respectively. A minimum of 50 cells were quantitated per condition. Statistical analysis was performed with ANOVA multiple comparison to wild-type condition with Dunett’s correction.

## Discussion

We have characterized a previously undescribed Rab5-independent, Rab14-dependent endosomal pathway. In this pathway, EEA1 is recruited to early endosomes in the absence of Rab5 (Figures 1, 3 and S3) and endosomal maturation requires Rab14 (Figures 3 and S5). This pathway appears to differ from previously described endocytosis already at the early stages of vesicle formation (Figure 2). We showed that synthetic molecules as CPP, as well as physiological molecules such as polyamines take advantage of the Rab5-independent, Rab14-dependent pathway to enter cells and reach lysosomes.

The molecules that we have found to be endocytosed via the Rab14-dependent pathway are characterized by their strong cationic nature. As Rab14 is expressed in most tissues (according to Protein Atlas and CCLE database) and seems to be present in the last eukaryote common ancestor ^97, 120^, it is possible that Rab14 is involved in a primordial endosomal pathway taken by cationic cargos. Whether this pathway is used by other types of cargos can now be assessed functionally in cells in which Rab14 is inactivated or invalidated.

Besides Rab14, no other Rab proteins could be evidenced to play a role in maturation of the endocytic pathway taken by CPPs, HDs, or polyamines. In particular, we did not find a Rab7 homolog that would be required for the acquisition of the Lamp1 marker in this pathway. Either Rab14 is the sole Rab necessary for cargo progression along the endocytic pathway used by CPPs, HDs, or polyamines or there is an as yet unknown Rab isoform that is redundant with Rab7 in this pathway. Because there is only marginal colocalization between Rab14 and Lamp1 ^93, 96^ (Figure S4), the possibility that Rab14 plays a dual role in the recruitment of EEA1 and Lamp1 is unlikely. Based on phylogeny and clustering analysis ^95, 97, 120^, Rab7 is most closely related to Rab9. However, cells lacking both Rab7 and Rab9 were not compromised in their ability to move cationic cargos along the Rab14-dependent pathway all the way down to Lamp1-containing vesicles (Figure 3). The Rab isoform that is redundant with Rab7, if it exists, remains therefore to be discovered.

There was minimal colocalization between CPP-, HP- or polyamine-containing endosomes and the Rab5 and Rab7 endosomal markers. This marginal entry could correspond to non-selective bulk liquid uptake, as it occurs during macropinocytosis^121^. Alternatively, a small fraction of these cargos may enter the Rab5-dependent endosomal pathway as previously reported for CPPs ^16, 18–23^.

There have been earlier circumstantial indications of the existence of a Rab5-independent pathway specifically in the context of viral infections, as well as trafficking to yeast vacuole ^122, 123^. It has been shown that some viruses use unconventional endocytosis, such as Herpes Simplex Virus 1 ^124^, SARS ^125^, Lassa virus, the Amr53b and We54 strains of LCMV (lymphocytic choriomeningitis virus) ^126–129^, Lujo virus ^130, 131^ and some influenza A strains ^132^ that appear to skip the early Rab5-positive endosomes. Rab14 also appears to be involved in viral trafficking, more specifically in the endocytosis of Ebola virus matrix protein VP40 ^133^. Additionally, Rab14 depletion delayed *Candida albicans*-containing phagosome maturation to Lamp1, even though Rab5 and Rab7 markers were still present during the maturation process ^134^. Possibly, the Rab5-independent, Rab14-dependent endosomal pathway is used by some pathogens to infect cells. Therefore, our findings may not only concern the physiological delivery of cationic cargo into cells, but could also be relevant for the search of anti-viral or antimicrobial drugs.

## Author contributions

Conception and design of study: ET and CW Acquisition of data: ET

Analysis and/or interpretation of data: ET, YH, MF and CW Funding acquisition: YH, MF and CW

Resources: MF and CW

Drafting the manuscript: ET and CW

Revising the manuscript and approval of the submitted version: all authors

## Acknowledgements

The laboratory of YH is supported by Grant-in-Aid for Young Scientists from the Ministry of Education, Culture, Sports, Science and Technology (MEXT) of Japan (grant number 20K15739). The laboratory of MF is supported by Grant-in-Aid for Scientific Research(B) from the MEXT (grant number 19H03220) and Japan Science and Technology Agency (JST) CREST (grant number JPMJCR17H4). We are thankful to the Cellular Imaging Facility and Bioinformatics Competence Centre at the University of Lausanne for the resources provided and their technical help.

## Materials & Correspondence

Correspondence and requests for materials should be addressed to CW.

## MATERIAL AVAILABILITY

Further information and requests for resources and reagents should be directed to and will be fulfilled by Christian Widmann. Plasmids generated in this study will be deposited to Addgene. This study did not generate any new unique reagents.

## EXPERIMENTAL MODEL AND SUBJECT DETAILS

### Cell lines

All cell lines were cultured in 5% CO2 at 37°C. HeLa (ATCC: CCL-2) cells were cultured in RPMI media (Thermo Fisher, Cat# 61870044) supplemented with 10% heat-inactivated fetal bovine serum (FBS; Thermo Fisher, Cat# 10270-106). MDCK-II parental and knock-out cell lines^99^ (available from RIKEN BioResource Research Center Cell Bank (https://cell.brc.riken.jp/en); Cat#: RCB5099–RCB5148) were cultured in DMEM (Thermo Fisher, Cat# 10566016). MDCK-II Rab5 knock-out and degron-Rab5A-expressing cell line will be described elsewhere (Homma et al., manuscript in preparation).

## REAGENTS

### Chemicals

Live Hoechst 33342 (Sigma, Cat# CDS023389) was aliquoted and stored at -20°C. AlexaFluor488-, AlexaFluor568-, AlexaFluor647-labeled human transferrin was dissolved in PBS at 5 mg/ml and stored at 4°C (Thermo Fisher, Cat# T13342, T23365 and T23366). TMR-labelled 10,000 neutral dextran was dissolved in PBS at 10 mg/ml and stored at -20°C (Thermo Fisher, Cat# D1816). AlexaFluor647-labeled and biotinylated epidermal growth factor (EGF) was dissolved in water at 1 mg/ml, aliquoted and stored at -20°C (Thermo Fisher Cat# E35351). EIPA (stock concentration 10 mM), IPA3 (stock concentration 5 mM), ML7 (stock concentration 10 mM), CytoD (stock concentration 1 mM), Jas (stock concentration 1 mM) a kind gift from Stefan Kunz laboratory, were dissolved in DMSO, aliquoted and stored at -20°C. LY294002 was dissolved in DMSO at 20 mg/ml, aliquoted and stored at -20°C (Sigma Aldrich, Cat# 440202). Wortmannin was aliquoted and stored at -20°C (Sigma Aldrich, Cat# W1344). Indole-3-acetic acid (IAA) (Sigma-Aldrich, Cat# I2886) was dissolved in ethanol, aliquoted and stored at -20°C. Doxycycline (Sigma-Aldrich, Cat# D3447) was dissolved in DMSO, aliquoted and stored at -20°C.

### Antibodies

The mouse monoclonal anti-EEA1, stored at -20°C (BD Transduction Laboratories, Cat# 610457) and anti-Lamp1, stored at 4°C (BD Pharmingen, Cat# 555798) antibodies were used in immunofluorescence experiments. Donkey polyclonal anti-mouse Cy3 secondary antibody was aliquoted and stored at -20°C in glycerol (Jackson ImmunoResearch, Cat# 715-165-150). Phospho-AKT (Ser473) rabbit polyclonal antibody was stored at -20°C (Cell signaling, Cat# 92715) and was used for western blotting. Anti-Rab5C antiserum, which can recognize all three Rab5 isoforms ^135^, was stored at -20°C.

## METHOD DETAILS

### Confocal microscopy

Confocal microscopy experiments were done on live cells. Cells were seeded onto glass bottom culture dishes (MatTek, corporation Cat# P35G-1.5-14-C) and treated as described in the Figures. For nuclear staining, 10 μg/ml live Hoechst 33342 (Molecular probes, Cat# H21492) was added in the culture medium 5 minutes before washing cells twice with media. After washing, cells were examined at the indicated time point with a plan Apochromat 63x oil immersion objective mounted on a Zeiss LSM 780 laser scanning fluorescence confocal microscope equipped with gallium arsenide phosphide detectors and three lasers (a 405 nm diode laser, a 458-476-488-514 nm argon laser, and a 561 nm diode-pumped solid-state laser). Cell images were acquired at a focal plane near the middle of the cell making sure that nuclei were visible. Experiments at 20°C were done using an incubation chamber set at 20°C, 5% CO2 and visualized with a Zeiss LSM710 Quasar laser scanning fluorescence confocal microscope equipped with either Neofluar 63x, 1.2 numerical aperture (NA) or plan Neofluar 100x, 1.3 NA plan oil immersion objective (and the same lasers as above).

### Colocalization

Colocalization assessment between endocytosed material and a given endosomal marker was performed on confocal images by visual assessment, switching back and forth between the color channels. The samples were randomized to blind the experimentators from the nature of the samples they were analyzing. The randomization script is available at https://github.com/BICC-UNIL-EPFL/randomizer.

The visual quantitation was validated by Mander’s coefficient calculation performed on the same samples using the JaCoP plugin in ImageJ. Examples of colocalization quantitation analysis is shown in Figure S1A.

### Immunofluorescence

Immunofluorescence experiments for the localization of endogenous and ectopically expressed EEA1 and Lamp1 endosomal markers was performed as described ^57^.

Briefly, cells were plated on poly L-lysine-coated coverslips and fixed with 4% paraformaldehyde for 20 minutes at room temperature at the indicated time points after treatment. Following a 5-minute permeabilization at room temperature in PBS, 0.25% triton X100, the samples were blocked for 20 minutes at room temperature in PBS, 3% BSA. Incubation with primary antibodies was done for two hours at room temperature in PBS, 1% BSA. The cells were then incubated for 45 minutes at room temperature with Cy3-labelled secondary antibodies in the same buffers as above. Coverslips were finally incubated 5 minutes with PBS, 10 μg/ml Hoechst. Three PBS washes were done between each incubation steps. Coverslips were mounted in Fluoromount-G (cBiosience, Cat# 00-4958-02). Samples were visualized with a Zeiss LSM780 confocal microscope.

### Transient transfection

Calcium phosphate based transfection of HeLa cells was performed as previously described ^136^. Briefly, cells were plated overnight in DMEM (Invitrogen, Cat# 61965) medium supplemented with 10% heat-inactivated FBS (Invitrogen, Cat# 10270-106),

2.5 μg of total plasmid DNA of interest was diluted in water, CaCl2 was added and the mixture was incubated in presence of HEPES 2x for 60 seconds before adding the total mixture drop by drop to the cells. Media was changed 10 hours after. Transient transfection in MDCK cells was done with Lipofectamin 2000 reagent according to supplier’s instructions (Thermo Fisher, Cat# 11668030).

### PI3-kinase inhibitors

For the colocalization experiments, cells ectopically expressing GFP-EEA1, were preincubated in the presence or in the absence of PI3K inhibitors 25 μM LY294002 or 10 μM wortmannin for 30 minutes, then Alexa548-transferrin or TMR-CPP were added to the cells for 5 minutes. Cells were washed on ice with RPMI, 10% FBS and incubated in same media in the presence of the inhibitors. Cells were visualized with LSM780 confocal microscope at the indicated time points. Colocalization between fluorescent cargo and EEA1 was visually quantitated. For western blotting experiments phosphorylated AKT was used as a proxy of PI3K activity. To stimulate phosphorylation, cells were incubated in serum-free medium for one hour. Medium was then changed to RPMI, 10% serum and cells were incubated for 20 minutes with 25 μM LY294002. Phosphorylated AKT was detected using rabbit anti-phosphoAKT antibody (Cell signaling, Cat# 92715).

### Macropinocytosis inhibition

Cells ectopically expressing GFP-EEA1 were starved overnight to stimulate macropinocytosis. Media was then changed to RMPI containing 10% FBS and cells were preincubated for 30 minutes with the indicated macropinocytosis inhibitors (kind gift from Dr. Stephan Kunz lab). Cells were pulsed for 5 minutes with TexasRed-dextran or TMR-TAT-RasGAP317-326, washed and visualized under confocal microscope in RPMI, 10% FBS in the presence of macropinocytosis inhibitors.

### Plasmid constructs

The RFP-hRab5A.dn3 (#921) plasmid encoding RFP-labeled version of human Rab5A protein was from Addgene (Cat# 14437). The mCh-hRab7A (#922) plasmid encoding mCherry-labeled version of human Rab7A protein was from Addgene (Cat# 61804). GFP-hRab5A.dn3 (#966), GFP-hRab7A.dn3 (#968), GFP-DynI.dn3 (#963) plasmid encoding GFP-labeled versions of the indicated human proteins, as well as dominant negative isoforms of the following proteins GFP-hRab5A(S34N).dn3 (#961), GFP-Rab7A(T22N).dn3 (#969), GFP-DynI(K44A).dn3 (#964) were a kind gift from Stefan Kunz laboratory. mCh-hRab5(S34N) (#933), plasmid encoding a mCherry-labeled dominant negative mutant version of human Rab5A was from Addgene (Cat# 35139). GFP-hRab5B.dn3 (#1008) plasmid encoding GFP-labeled wild-type version of human Rab5B isoform was from Addgene (Cat# 61802). GFP-hRab5B(S34N) (#1067) plasmid encoding GFP-labeled dominant negative mutant version of human Rab5B isoform was introduced to plasmid #1008 using Q5 Site-Directed Mutagenisis kit (NEB, Cat# E0554S) according to manufacturer’s instructions using forward primer #1554 (AGTGGGAAAGaacAGCCTGGTATTAC) and reverse primer #1555 (GCAGATTCTCCCAGCAGG). GFP-Rab5C.dn3 (#1074) plasmid encoding GFP-labeled version of human Rab5C isoform was from Genescript (Cat# OHu09753C). CFP-hRab5C(S35N).dn3 (#1006) encoding Cerulean-labeled dominant negative mutant version of human Rab5C isoform was from Addgene (Cat# 11504). GFP-hEEA1 (#970) and hLamp1-GFP.dn3 (#971) encoding GFP-labeled version of human EEA1 and Lamp1, respectively were from Addgene (Cat# 42307 and 34831). BFP-EEA1 (#1009) plasmid was generated by subcloning GFP-hEEA1 (#970) into a BFP-hRab7A-Myc backbone (#1005, Addgene Cat# 79803) through ligation of both plasmids after digestion with BamHI (NEB, Cat# R313614) and BspEI (NEB, Cat# R0540S). hLamp1-BFP (#1016) plasmid encoding BFP-labeled version of human Lamp1 protein was from Addgene (Cat# 98828). GFP-hRab14.dn3 (#1017), GFP-hRab21.dn3 (#1023), GFP-hRab22.dn3 (#1018) and GFP-hRab31.dn3 (#1019) plasmids encoding GFP-labeled versions of the indicated human wild-type proteins were from Addgene (Cat# 49549, 83421, 49600 and 49610, respectively). GFP-hRab14(S25N).dn3 (#1037) and GFP-hRab14(N124I).dn3 (#1038) plasmids encoding

GFP-labeled versions of dominant negative Rab14 mutant were from Addgene (Cat#49594 and 49593). GFP-hRab21(T31N) (#1039) plasmid encoding GFP-labeled version of dominant negative Rab21 mutant was from Addgene (Cat# 83423). GFP-hRab22(S19N) (#1068) was generated using Q5 Site-Directed Mutagenisis kit (NEB, Cat# E0554S) according to manufacturer’s instructions with forward primer #1556 (TGTAGGTAAAaacAGTATTGTGTGGCGG) and reverse primer #1557 (CCTGTATCCCCGAGCAGA) on plasmid #1018 that encodes the wild-type isoform of human Rab22 protein.

### Peptides

TAT-RasGAP317-326 is a retro-inverso peptide (i.e. synthesized with D-amino-acids in the opposite direction compared to the natural sequence) labeled or not with FITC or TMR. The TAT moiety corresponds to amino-acids 48–57 of the HIV TAT protein (RRRQRRKKRG) and the RasGAP317–326 moiety corresponds to amino-acids from 317 to 326 of the human RasGAP protein (DTRLNTVWMW). These two moieties are separated by two glycine linker residues in the TAT-Ras-GAP317–326 peptide. FITC-or TMR-bound peptides without cargo: TAT, MAP (KLALKLALKALKAALKLA), Penetratin (RQIKWFQNRRMKWKK), Transportan (GWTLNSAGYLLGKINLKALAALAKKIL), R9 (RRRRRRRRR), were synthesized in retro-inverso conformation. FITC-labeled homeodomain: OTX2-HD (QRRERTTFTRAQLDVLEALFAKTRYPDIFMREEVALKINLPESRVQVWFKNRRAKCRQQQ). All peptides were synthesized by SBS Genetech, China and resuspended to 1 mM in water.

### Polyamine labeling

Spermine fluorescent labeling with CF405M or CF594 dyes was performed using a Mix-n-Stain Small Ligand Labeling Kit (Biotium, Cat# 92362 and 92352, respectively) according to manufacturer’s instructions. The labeling efficiency was assessed by reverse-phase HPLC at Protein and Peptide Chemistry Facility at University of Lausanne.

### Statistical analysis

Statistical analysis was performed on non-normalized data, using GraphPad Prism 7. All measurements were from biological replicates. Unless otherwise stated, the vertical bars in the graph represent the standard deviation of mean from at least three independent experiments.

### Data availability

Image randomization script can be found at: https://github.com/BICC-UNIL-EPFL/randomizer.

---

**Figure S1.**
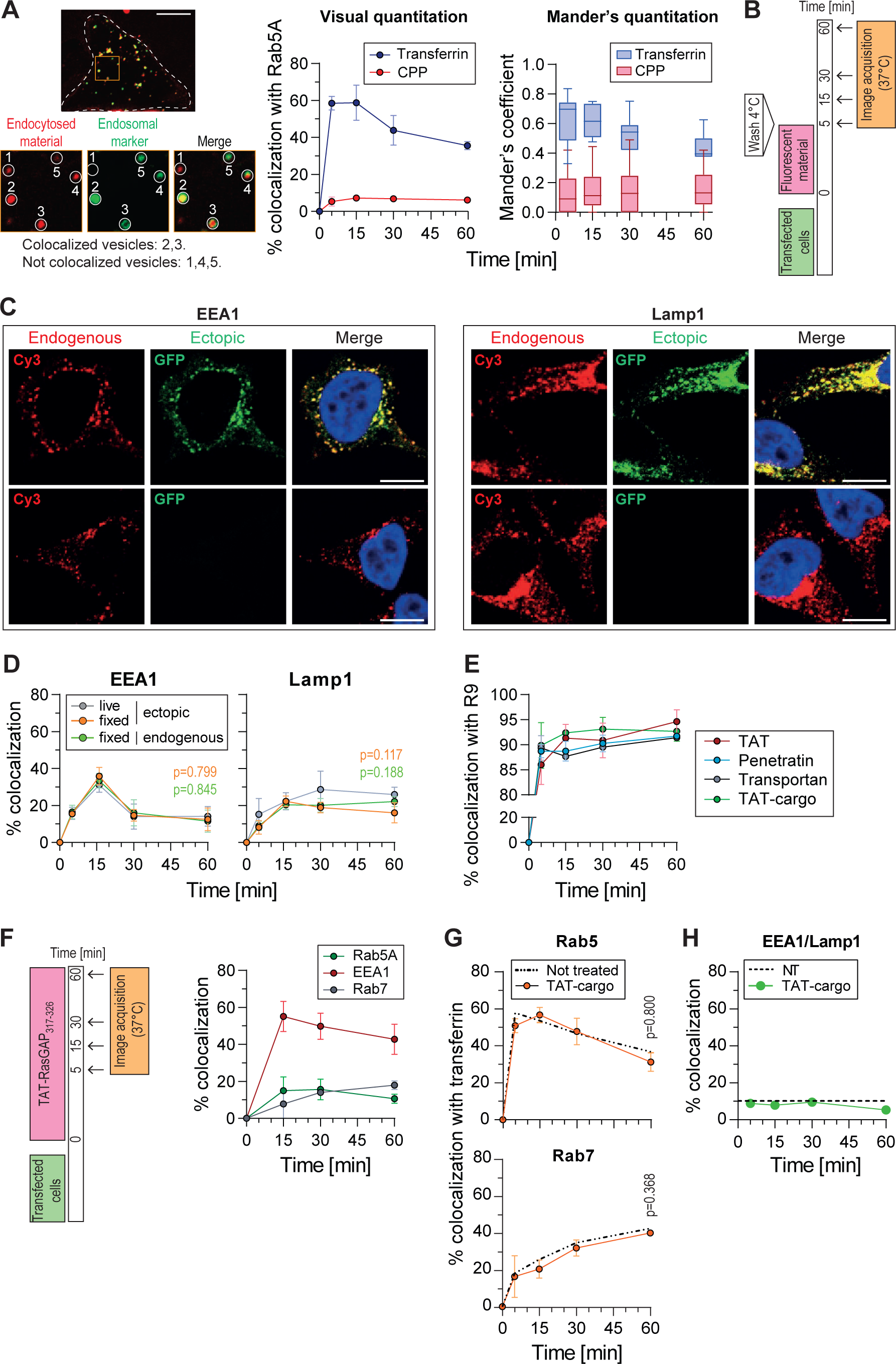
Colocalization quantitation and experimental setup. **(A)** Colocalization quantitation between fluorescently labeled endosomal material and endocytic markers. Left: representative confocal images of HeLa KCNN4 knock-out cells expressing GFP-Rab5A, in the presence of 20 ug/ml AlexaFluor568-Transferrin for 5 minutes. Colocalization assessment between endocytosed material and a given endosomal marker was performed on confocal images by visual assessment, switching back and forth between the color channels. The samples were randomized to blind the experimentators from the nature of the samples they were analyzing. The images shown on the left are cropped regions of cell transfected with GFP-Rab5 and incubated with AlexaFluor568-transferrin. The circles numbered 1 to 5 depict examples of colocalization versus non-colocalization between Rab5 and transferrin. Scale bar: 10 μm. Graphs on the right-hand side: our visual quantitation (mean ± SD of three independent experiments performed on 165 cells per condition) was validated by Mander’s coefficient calculation performed on the same samples using the JaCoP plugin in ImageJ (shown as box plots). **(B)** Scheme of the pulse chase experiments used in the experiments depicted in the figures. Cells were incubated five minutes with various fluorescent material, washed, and the endosomal maturation followed overtime. **(C)** Ectopic expression does not alter the subcellular location of endosomal markers. Ectopic (top row) and endogenous (bottom row) location of EEA1 and Lamp1. Scale bar: 10 μm. **(D)** Quantitation of the colocalization between ectopic or endogenous EEA1 or Lamp1 with Alexa568-transferrin in live or fixed KCNN4 knock-out HeLa cells. The data correspond to mean ± SD of three independent experiments. The p-values were calculated using ANOVA analysis with Dunett’s correction based on AUC values from fixed samples compared to live samples. **(E)** Colocalization quantitation of the indicated FITC-CPPs with TMR-R9. The data correspond to the mean ± SD of three independent experiments. **(F)** Colocalization between TAT-RasGAP317-326 (40 μM) and GFP-tagged Rab5, EEA1 or Rab7 in HeLa KCNN4 knock-out cell line. Cells were incubated in the continuous presence of TAT-RasGAP317-326. The data correspond to mean ± SD of three independent experiments. **(G)** Colocalization between transferrin and Rab5 (top) or Rab7 (bottom) in the presence or in the absence of 40 μM TAT-RasGAP317-326. The data correspond to mean ± SD of three independent experiments. The p-values were calculated based on AUC analysis. **(H)** Colocalization quantitation between ectopically expressed BFP-EEA1 and Lamp1-GFP in the presence or in the absence of 40 μM TAT-RasGAP317-326. The data correspond to mean ± SD of three independent experiments.

**Figure S2.**
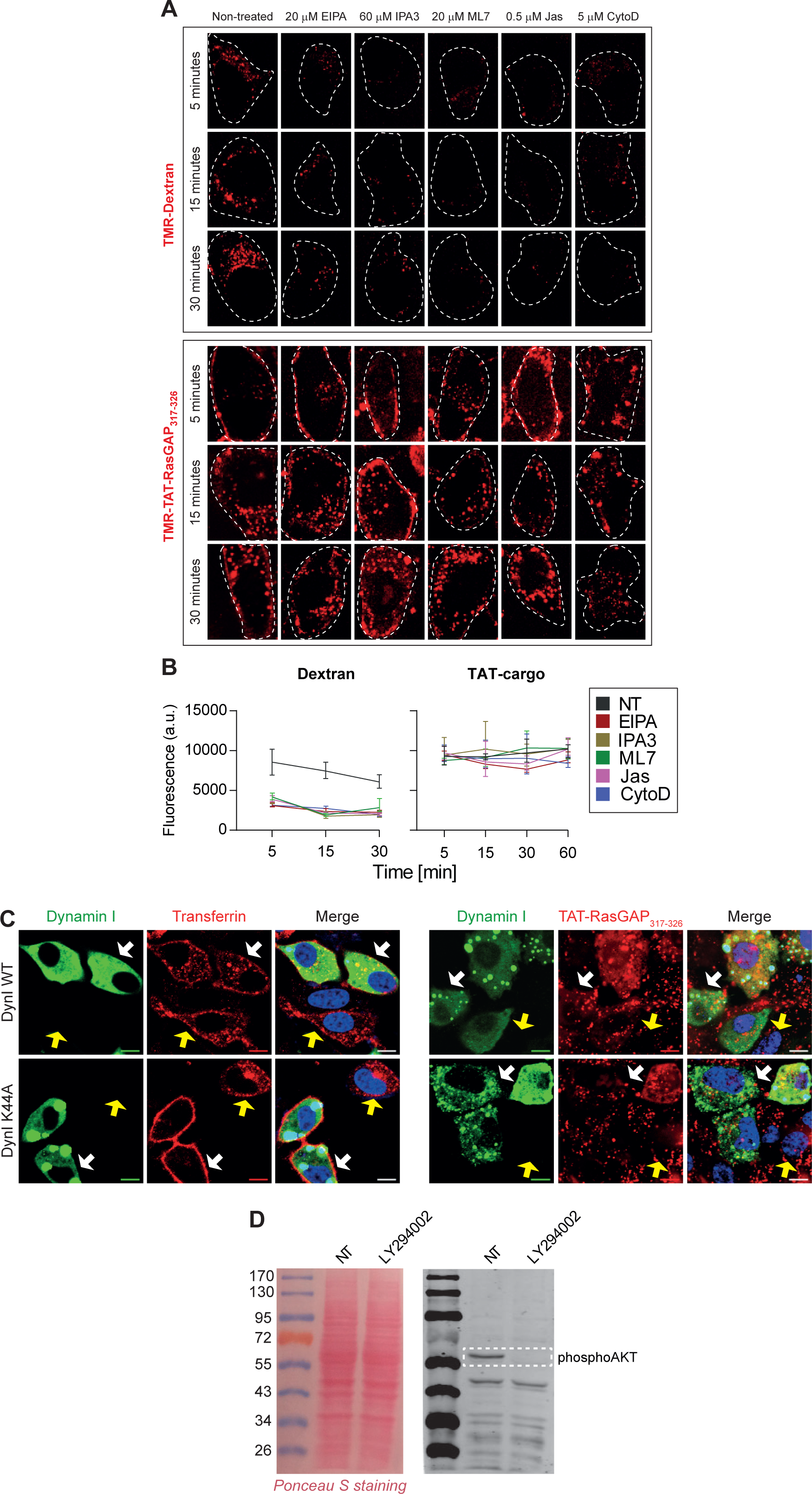
Endosomal vesicle formation inhibitors do not affect CPP uptake. **(A)** Representative confocal images of HeLa KCNN4 knock-out cells preincubated or not with the indicated inhibitors for 30 minutes prior to the addition of 0.2 mg/ml TMR-Dextran 10 kDa or 40 μM TMR-TAT-RasGAP317-326 for a 5-minute pulse. Confocal images were acquired at 5, 30 and 60 minutes after the addition of endosomal material. The inhibitors were present throughout the full duration of the experiment. **(B)** Total cell fluorescence quantitation based on images acquired in panel A. Results correspond to mean ± SD of three independent experiments (n>150 cells per condition). **(C)** Representative confocal images of HeLa KCNN4 knock-out cells ectopically expressing wild-type (DynI WT) or dominant negative (DynI K44A) version of dynamin I. Cells were incubated with 20 μg/ml AlexaFluor568-transferrin or 40 μM TAT-RasGAP317-326. Cell nuclei were labeled with live Hoechst. White arrows point to cells positively transfected with GFP-dynamin constructs and yellow arrows indicate non-transfected cells. Images were acquired at 15 minutes post incubation with the cargos. **(D)** Left panel: red Ponceau S stained membrane, right panel: phospho-AKT antibody signal detection. HeLa KCNN4 knock-out cells incubated in the presence or in the absence of 25 μM pan-PI3-kinase inhibitor, LY294002. To stimulate AKT phosphorylation cells were preincubated in a serum-free media for one hour. Phosphorylated AKT was used as a proxy for PI3-kinase activity.

**Figure S3.**
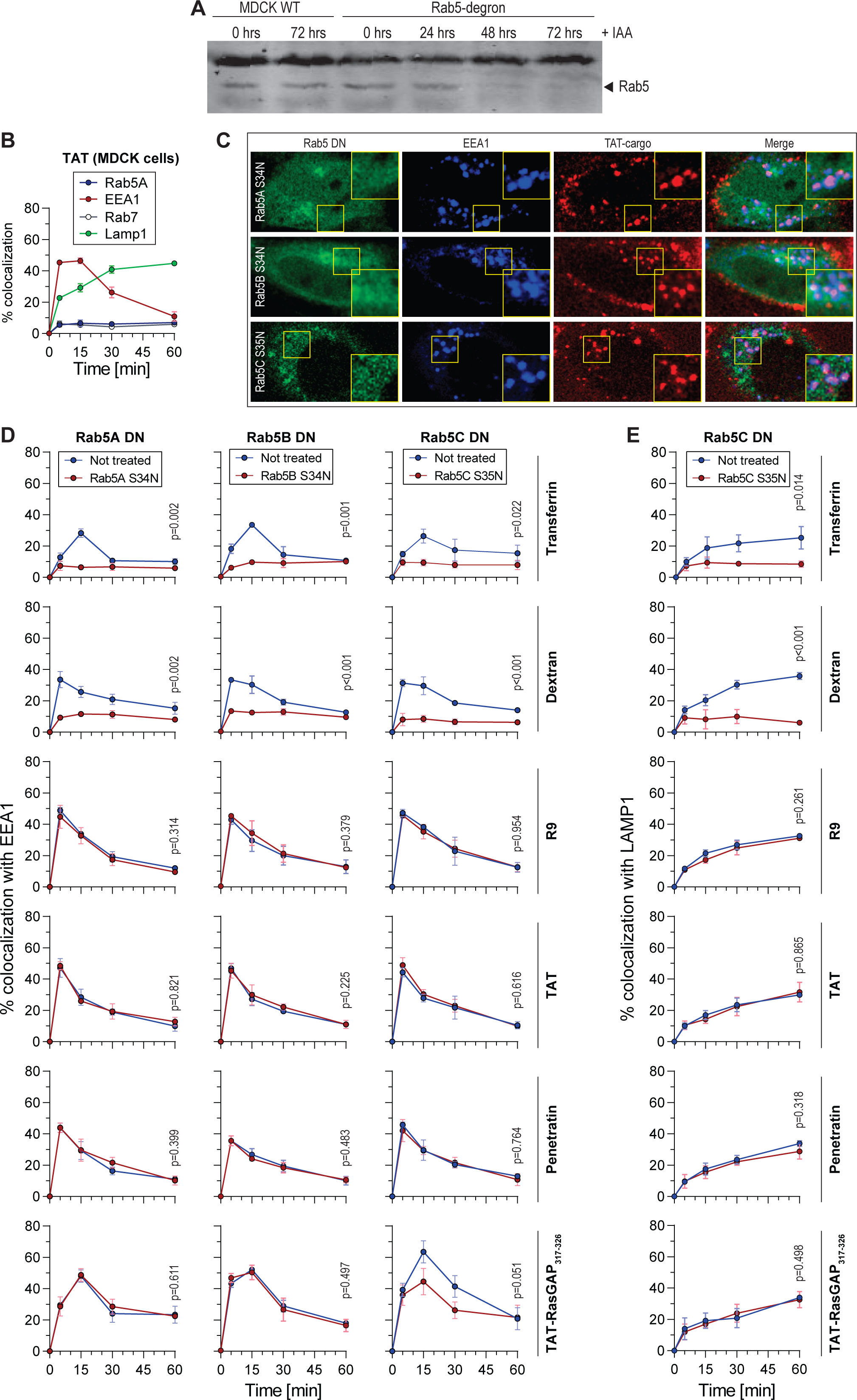
CPP endocytosis is Rab5-independent. **(A)** Western blot-mediated Rab5 detection in MDCK-II wild-type or Rab5-degron cell lysates incubated in the presence of 1 μg/ml doxycycline and 500 μM IAA for the indicated periods of time to induce degradation of Rab5A. **(B)** Colocalization of TAT-containing vesicles with the indicated endosomal markers. MDCK-II wild-type cells were incubated with 40 μM TMR-TAT for 5-minute pulse. **(C)** Representative confocal images of cells analyzed in panel D-E in HeLa KCNN4 knockout cells. **(D-E)** Colocalization quantitation of transferrin, dextran, and the indicated CPP with EEA1 (panel D) or Lamp1 (panel E) in cells transfected with the indicated Rab5 dominant negative constructs in HeLa KCNN4 knockout cells in a pulse experiment setting (see Figure S1B). The data correspond to mean ± SD of three independent experiments. Statistical analysis was performed using two-tailed t-test based on AUC values. The p-values correspond to the comparison between the cells within the same population transfected or not with the dominant negative constructs.

**Figure S4.**
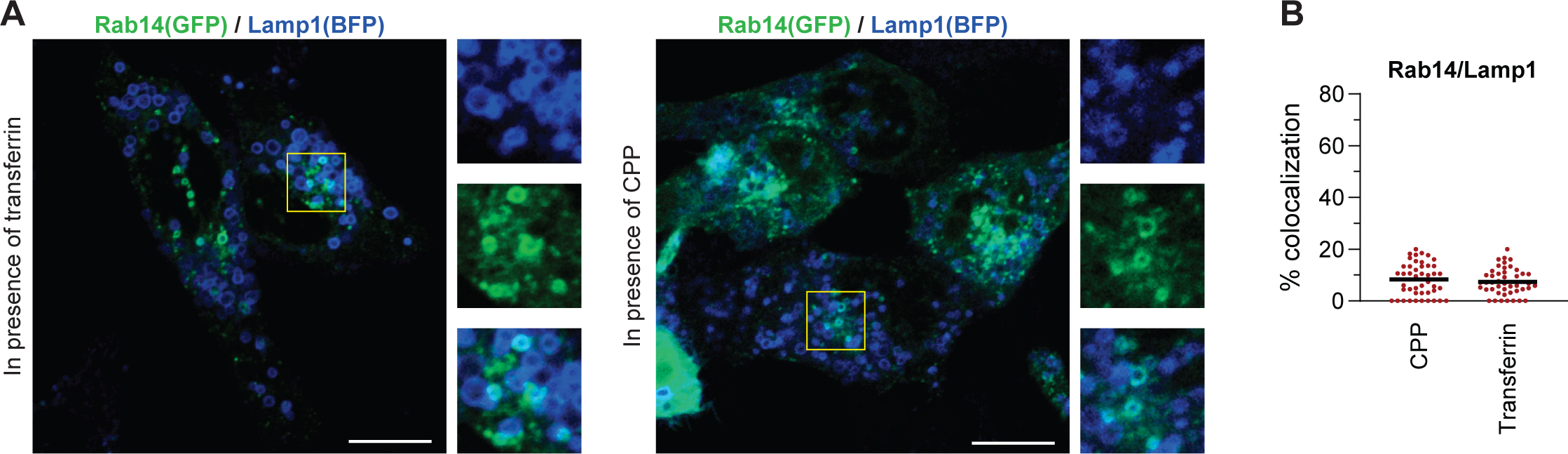
Limited colocalization between Rab14 and Lamp1. **(A)** Representative confocal images of HeLa KCNN4 knockout cells ectopically expressing GFP-Rab14 and Lamp1-BFP that were incubated for 5 minutes with 20 μg/ml AlexaFluor647-Transferrin or 40 μM TMR-TAT. Images were acquired at 30 minutes post-incubation with endosomal cargo. Scale bar: 10 μm. **(B)** Colocalization quantitation of Rab14 with Lamp1 from the experiment in panel A. A minimum of 50 cells were quantitated per condition.

**Figure S5.**
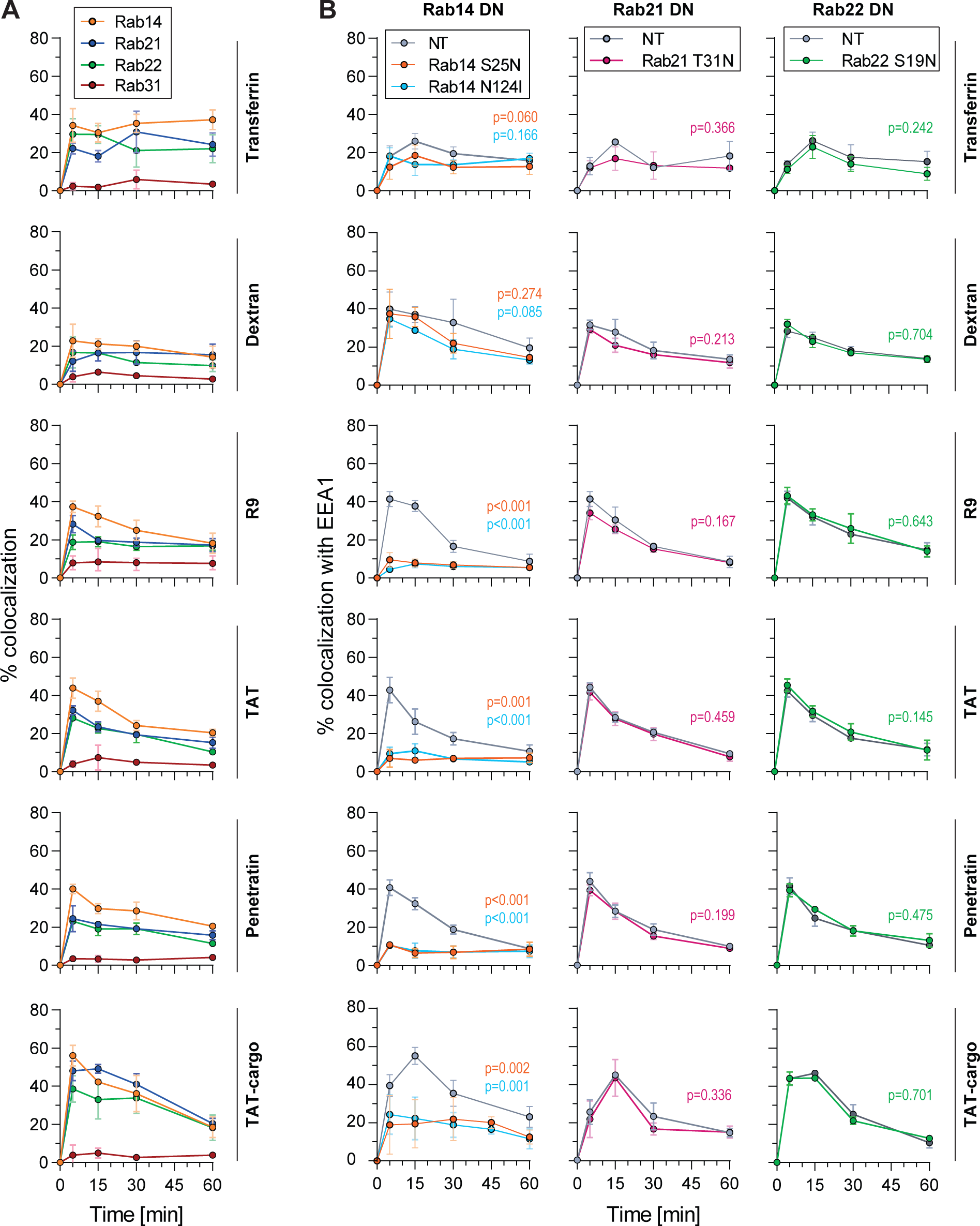
Rab14 dominant negative mutants block the maturation of CPP-containing endosomes. **(A)** Colocalization quantitation between the indicated endocytosed material and the indicated GFP-labeled Rab proteins ectopically expressed in HeLa KCNN4 knockout cells. The results correspond to mean ± SD of three independent experiments. **(B)** Colocalization quantitation between the indicated endocytosed material and EEA1 in cells transfected with Rab14, Rab21 or Rab22 dominant negative constructs. The data correspond to the mean ± SD of three independent experiments. Statistical analysis was performed using ANOVA test with Dunett’s correction, based on AUC values. The p-values correspond to the comparison between the cells within the same population transfected or not with the dominant negative constructs.

**Figure S6.**
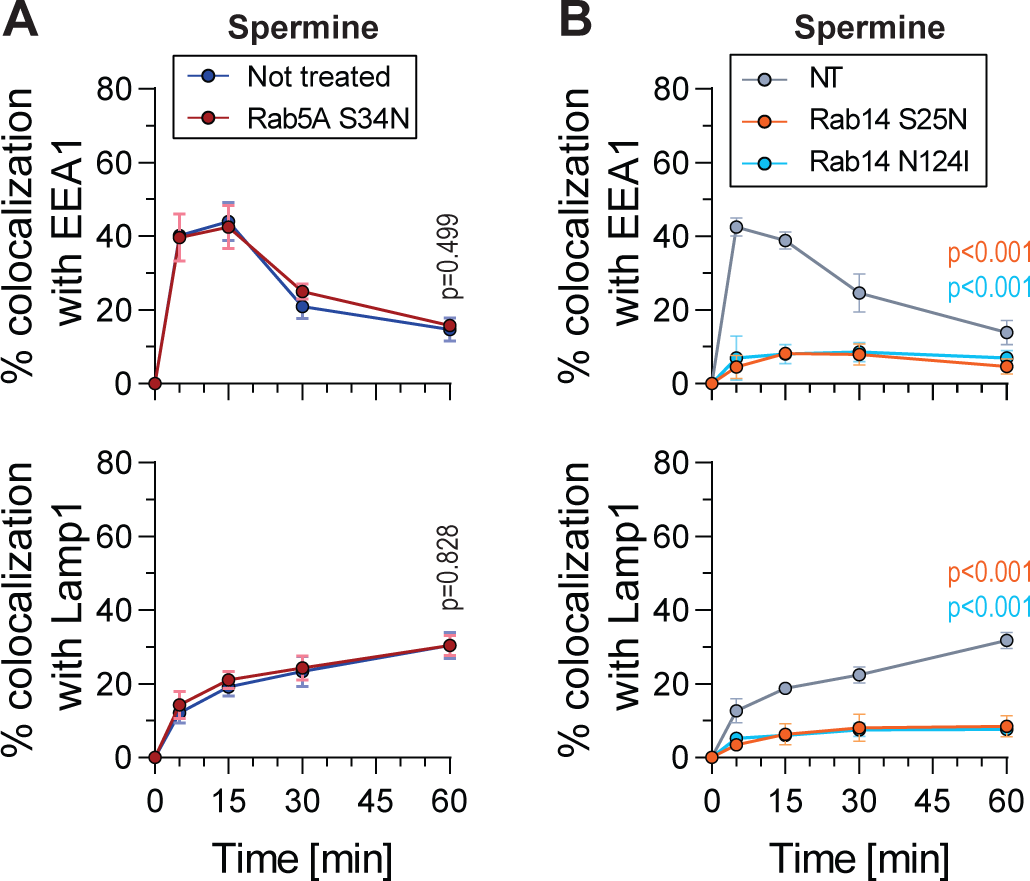
Spermine follows a Rab5-independent, Rab14-dependent endosomal maturation. **(A)** Colocalization between 5 μM spermine with EEA1 (top) or Lamp1 (bottom) in the Rab5A dominant negative background. HeLa KCNN4 knockout cells were incubated for 5-minute pulse with endosomal cargo. The results correspond to mean ± SD of three independent experiments. Statistical analysis was performed using two-tailed t-test based on AUC values. The p-values correspond to the comparison between the cells within the same population transfected or not with the dominant negative constructs. **(B)** Colocalization between 5 μM spermine with EEA1 (top) or Lamp1 (bottom) in HeLa KCNN4 knockout cells expressing Rab14 dominant negative mutants. The cells were exposed to the cargos during a 5-minute pulse. The results correspond to mean ± SD of three independent experiments. Statistical analysis was performed using ANOVA test with Dunett’s correction, based on AUC values. The p-values correspond to the comparison between the cells within the same population transfected or not with the dominant negative constructs.

